# Cerebral μ-opioid and CB_1_-receptor systems have distinct roles in human feeding behavior

**DOI:** 10.1101/2020.12.17.423284

**Authors:** Tatu Kantonen, Tomi Karjalainen, Laura Pekkarinen, Janne Isojärvi, Kari Kalliokoski, Valtteri Kaasinen, Jussi Hirvonen, Pirjo Nuutila, Lauri Nummenmaa

## Abstract

Eating behavior varies greatly between healthy individuals, but the neurobiological basis of these trait-like differences in feeding remains unknown. Central µ-opioid receptors (MOR) and cannabinoid CB_1_-receptors (CB_1_R) regulate energy balance via multiple neural pathways, promoting food intake and reward. Because obesity and eating disorders have been associated with alterations in the brain’s opioid and endocannabinoid signaling, the variation in MOR and CB_1_R system function could potentially underlie distinct eating behavior phenotypes. In this retrospective positron emission tomography (PET) study, we analyzed [^11^C]carfentanil PET scans of MORs from 92 healthy subjects (70 males and 22 females), and [^18^F]FMPEP-*d*_*2*_ scans of CB_1_Rs from 35 subjects (all males, all also included in the [^11^C]carfentanil sample). Eating styles were measured with the Dutch Eating Behavior Questionnaire (DEBQ). We found that lower cerebral MOR availability was associated with increased external eating – individuals with low MORs reported being more likely to eat in response to environment’s palatable food cues. CB_1_R availability was associated with multiple eating behavior traits. We conclude that although MORs and CB_1_Rs overlap anatomically and functionally in the brain, they have distinct roles in mediating individual feeding patterns.

## 1. Introduction

Obesity is one of the leading public health issues, resulting from individuals’ long-term excessive energy intake in relation to energy expenditure (Guyenet and Schwartz, 2012). Yet, humans vary greatly in their choices and habits related to food intake quantity and quality i.e. eating behavior (French et al., 2012; Larson and Story, 2009). Trait-like eating behaviors have been associated with multiple clinical eating disorders in addition to obesity (Baños et al., 2014; Barrada et al., 2016; Cebolla et al., 2014; Wardle, 1987), but also non-obese individuals vary in how they control their feeding (Yeomans et al., 2008). Interacting with peripheral hormones, central nervous system (CNS) integrates hunger and satiety signals with environmental stimuli to regulate food intake (Guyenet and Schwartz, 2012). Large-scale genome-wide association studies have identified limbic system, hippocampus and hypothalamus to be key regions in the CNS contributing to individual’s body mass index (BMI) and eating behavior (Locke et al., 2015; Silventoinen and Konttinen, 2020). Central regulation of feeding is however constantly challenged by the modern environment characterized by abundance of palatable and energy-dense food products, promoting feeding independently of metabolic needs (Berthoud, 2012; Hill and Peters, 1998).

Palatability and hedonic properties of food are centrally mediated by µ-opioid receptor (MOR) system (Gosnell and Levine, 2009; Nummenmaa et al., 2018). Both endogenous and exogenous opioids stimulate feeding, especially via hedonic hotspots of nucleus accumbens, insula and frontal cortex (Castro and Berridge, 2017; Mena et al., 2011; Nogueiras et al., 2012; Smith and Berridge, 2007). Conversely, opioid antagonists reduce food intake and related hedonic responses in rodents (Nogueiras et al., 2012) and humans (Yeomans and Gray, 2002; Ziauddeen et al., 2013). Human positron emission tomography (PET) studies have revealed that obesity associates with decrease of MORs in appetite regulating brain areas (Burghardt et al., 2015; Karlsson et al., 2015), and insular MORs are lowered in patients with bulimia nervosa proportionally to fasting behavior (Bencherif et al., 2005). Central MOR system function varies considerably also in healthy humans (Kantonen et al., 2020), and traits linked with feeding control such as impulsivity are associated with MOR availability (Love et al., 2009). Nevertheless, the association between the MOR system and specific patterns of eating behavior remains elusive.

Feeding is also regulated by brain’s endocannabinoid system, which overlaps anatomically and functionally with the central MORs (Cota et al., 2006). The most abundant central cannabinoid receptors are the CB_1_-receptors (CB_1_Rs), which regulate food intake through circuits of ventral striatum, limbic system and hypothalamus (Bermudez-Silva et al., 2012; Mechoulam and Parker, 2013). Functional interplay between MOR and CB_1_R systems is highlighted in animal studies, where CB_1_R-antagonists and MOR-antagonists have synergistic effect on reducing food intake (Rowland et al., 2001), and CB_1_R-antagonist can be used to block MOR-agonist induced food intake and vice versa (Solinas and Goldberg, 2005). MOR-agonists also directly increase endocannabinoid concentration and CB_1_R-agonists increase opioid concentration in the brain, including nucleus accumbens (Solinas et al., 2004; Vigano et al., 2004). In humans, CB_1_R-antagonist rimonabant showed promise as an anti-obesity drug, but was withdrawn due to psychiatric side effects (Di Marzo and Després, 2009). More nuanced understanding of CB_1_R system and feeding is clearly required to enable further pharmacological advancement.

### 1.1. The current study

Accumulating evidence suggests that variation in central MOR and CB_1_R function could be linked to feeding and pathological eating behavior traits in humans, but it remains unresolved what facets of feeding they govern in humans. Individual differences in eating patterns can be conceptualized based on the psychological mechanisms that contribute to or attenuate development of overweight. In such conceptualization, *emotional eating* refers to reactive overeating to distress or negative emotions, while *external eating* refers to tendency to overeat in response to attractive food-cues. Finally, *restrained eating* refers to the tendency to eat less than desired (Van Strien et al., 1986b; Van Strien et al., 2009; Van Strien et al., 2012a). The emotional and external overeating are based on psychosomatic and externality theories of eating behavior, while restrained eating dimension centers around food intake self-inhibition (Van Strien et al., 1986a). Such consistent patterns contribute to differences in weight gain and maintenance (Van Strien et al., 2009; Van Strien et al., 2012b), and they can be measured using The Dutch Eating Behavior Questionnaire (DEBQ) (Van Strien et al., 1986a). Here we compiled 92 [^11^C]carfentanil PET scans targeting the MOR system and 35 [^18^F]FMPEP-*d_2_* PET scans of CB_1_R system and corresponding DEBQ data in a retrospective PET database study. We tested whether the MOR and CB_1_R availabilities in food-intake-regulating brain areas associate with individual eating behavior traits measured with DEBQ.

## 2. Materials and Methods

### 2.1 Subjects

The study subjects were historical controls retrieved from the AIVO neuroinformatics database (http://aivo.utu.fi), a large-scale molecular image database hosted by Turku PET Centre. We identified all the [^11^C]carfentanil and [^18^F]FMPEP-*d*_*2*_ baseline PET studies accompanied with completed DEBQ form (Van Strien et al., 1986a). Exclusion criteria were neurologic and psychiatric disorders, current use of medications that could affect CNS or abuse of alcohol or illicit drugs. Subjects were not preselected on the basis of weight or eating habits. Final sample consisted of 92 subjects (70 males and 22 females) scanned with [^11^C]carfentanil from five distinct projects with three different PET scanners. The [^18^F]FMPEP-*d*_*2*_ sample consisted of 35 males, all of which were also all included in the [^11^C]carfentanil male sample. All subjects of the [^18^F]FMPEP-*d*_*2*_ subsample were nonsmoking males, while in the [^11^C]carfentanil sample seven females smoked. All [^18^F]FMPEP-*d*_*2*_ scans were carried out with GE Discovery VCT PET/CT (GE Healthcare). The original data were acquired between 2011 and 2019. Characteristics of the study sample are summarized in **Table 1**, and the information of smoking status and PET scanners are detailed in **Supplementary Table 1**. The study was conducted in accordance with the Declaration of Helsinki and approved by the Turku University Hospital Clinical Research Services. The subjects had signed written informed consent forms and completed the DEBQ forms as a part of the original study protocols.

**Table 1.** Characteristics of the studied subjects.

|  | <sup>[11</sup>C]carfentanil scans |  |  |  |  |  |
| --- | --- | --- | --- | --- | --- | --- |
|  | Males (n = 70) |  |  | Females (n = 22) |  |  |
|  | mean | SD | range | mean | SD | range |
| Age (years) | 27.4 | 7.5 | 19–58 | 47.7 | 10.0 | 20–59 |
| BMI (kg/m<sup>2</sup>) | 24.5 | 2.8 | 19–31 | 23.7 | 3.1 | 18–31 |
| Total DEBQ score | 67.0 | 12.8 | 40–109 | 73.4 | 12.8 | 46–97 |
| Emotional eating score | 21.0 | 6.8 | 13–40 | 22.1 | 5.7 | 13–32 |
| External eating score | 24.7 | 6.5 | 10–43 | 25.0 | 5.4 | 13–34 |
| Restrained eating score | 21.3 | 5.5 | 11–39 | 26.2 | 5.5 | 10–33 |
| Injected activity (MBq) | 277.0 | 77.9 | 223–508 | 352.3 | 125.5 | 234–519 |

|  | <sup>[18</sup>F]FMPEP-d<sub>2</sub> scans |  |  |
| --- | --- | --- | --- |
|  | Males (n = 35) |  |  |
|  | mean | SD | range |
| Age (years) | 25.9 | 4.3 | 21–35 |
| BMI (kg/m<sup>2</sup>) | 24.5 | 3.1 | 19–31 |
| Total DEBQ score | 68.6 | 14.5 | 43–109 |
| Emotional eating score | 20.7 | 7.4 | 13–40 |
| External eating score | 27.1 | 6.0 | 14–43 |
| Restrained eating score | 20.8 | 5.6 | 12–32 |
| Injected activity (MBq) | 187.9 | 12.8 | 147–215 |

### 2.2. Eating behavior assessment with the DEBQ

The Dutch Eating Behavior Questionnaire (DEBQ) (Van Strien et al., 1986a) was used to quantify eating behavior. The DEBQ is a 33-item questionnaire with Likert-type scoring in each item (response options ranging from 1−5, from “Never” to “Very often”). It is divided in three dimensions measuring different behavioral traits: Emotional eating, External eating and Restrained eating (Van Strien et al., 1986b; Van Strien et al., 2009; Van Strien et al., 2012a). The DEBQ subscales have been designed to measure independent dimensions of feeding behavior (Van Strien et al., 1995), and the subscales have good internal consistency, dimensional validity and test-retest reliability (Cebolla et al., 2014; Malesza and Kaczmarek, 2019; Van Strien et al., 1986a; Wardle, 1987).

### 2.3. Image processing and modeling

PET images were preprocessed similarly using automated processing pipeline Magia (Karjalainen et al., 2020). [^11^C]carfentanil data preprocessing has been described previously (Kantonen et al., 2020). MOR availability was expressed as [^11^C]carfentanil binding potential (*BP*_ND_), which is the ratio of specifically bound radioligand to that of nondisplaceable radioligand in tissue (Innis et al., 2007). Occipital cortex served as the reference region (Frost et al., 1989). CB_1_R availability was expressed as [^18^F]FMPEP-*d*_*2*_ volume of distribution (*V*_T_), which was quantified using graphical analysis by Logan (Logan, 2000). The frames starting at 36 minutes and later since injection were used in the model fitting, since Logan plots became linear after 36 minutes (Logan, 2000). Plasma activities were corrected for plasma metabolites as described previously (Lahesmaa et al., 2018).

### 2.4. Statistical analysis

The study question was whether the DEBQ subscales (Emotional eating, External eating, Restrained eating) or Total DEBQ scores are associated with [^11^C]carfentanil *BP*_ND_ or [^18^F]FMPEP-*d*_*2*_ *V*_T_. We used Bayesian hierarchical modeling to analyze these effects in *a priori* regions of interest (ROIs). FreeSurfer (http://surfer.nmr.mgh.harvard.edu/) was used to extract 10 bilateral ROIs based on known regions with high density of MORs (Kantonen et al., 2020) and CB_1_Rs (Terry et al., 2010): amygdala, caudatus, cerebellum, dorsal anterior cingulate cortex, insula, middle temporal cortex, nucleus accumbens, orbitofrontal cortex, putamen, and thalamus. The Bayesian models were estimated using the R package brms (https://cran.r-project.org/web/packages/brms/index.html) that utilizes the Markov chain Monte Carlo sampling of RStan (https://mc-stan.org/users/interfaces/rstan). Because age influences [^11^C]carfentanil binding (Kantonen et al., 2020; Zubieta et al., 1999) and different PET scanners may yield slightly different *BP*_ND_ estimates (Nummenmaa et al., 2020), both age and PET scanner were controlled for in all [^11^C]carfentanil models. Age was also controlled for in all [^18^F]FMPEP-*d*_*2*_ *V*_T_ models (the scanner-adjustment was not needed since the [^18^F]FMPEP-*d*_*2*_ data were acquired using a single scanner). For both tracers, we created models separately for the Total DEBQ score as well as its subscales, adjusting for age. [^11^C]carfentanil *BP*_ND_ and [^18^F]FMPEP-*d*_*2*_ *V*_T_ were log-transformed to improve model fit (Kantonen et al., 2020). For both tracers, we estimated varying (random) intercepts for the subjects and ROIs, and varying (random) slopes across ROIs for the predictor of interest (e.g. Total DEBQ score) and age. Compared to a model where the regionally specific effects would be estimated using interaction term for ROI, the hierarchical model produces results that are partially pooled towards each other, thus accounting for the multiple comparisons performed (Gelman et al., 2012). For both tracers, we also estimated regionally varying (random) residual variances. For [^11^C]carfentanil data, we also estimated regionally varying (random) intercepts for the scanners. We used the standard normal distribution as a prior distribution for the regression coefficients of the predictors to provide regularization. The standard normal distribution was also used as the prior distribution for the standard deviation of the group-level (random) effects. Otherwise we used the default priors of brms. We used 1000 warmup samples, 1000 post-warmup samples and 10 chains, thus totaling 10 000 post-warmup samples. The sampling parameters were slightly modified to facilitate convergence (*adapt_delta* = 0.999; *max_treedepth* = 20). The samplings produced no divergent iterations and the Rhats were all 1.0, suggesting that the chains converged successfully.

To examine associations in the whole brain, we used nonparametric approach with SnPM13 (http://nisox.org/Software/SnPM13/) to create full-volume linear regression models for *BP*_ND_ and *V*_T_ values. We used p < 0.01 as the cluster-defining threshold, and only report clusters large enough to be statistically significant at FWE p < 0.05. 5000 permutations were used to estimate the null distribution. We created distinct models for Total DEBQ score and all the subscale scores, adjusting for age and also for PET scanner in [^11^C]carfentanil models. To rule out the possible effects of sex, smoking and BMI, we replicated the [^11^C]carfentanil full volume analysis with these additional covariates. The [^18^F]FMPEP-*d*_*2*_ models were also replicated with BMI as additional covariate (there were no smokers or females in the [^18^F]FMPEP-*d*_*2*_ data).

## 3. Results

Mean distribution of MORs and CB_1_Rs is shown in **Figure 1**. Correlations between the DEBQ subscales were positive but only modest, strongest being between Emotional and External eating (*r* = +0.33). BMI had a significant correlation only with Restrained eating (*r* = +0.27). Correlations with p-values are presented in **Supplementary Figure 1** and **Supplementary Table 2**.

**Figure 1.**
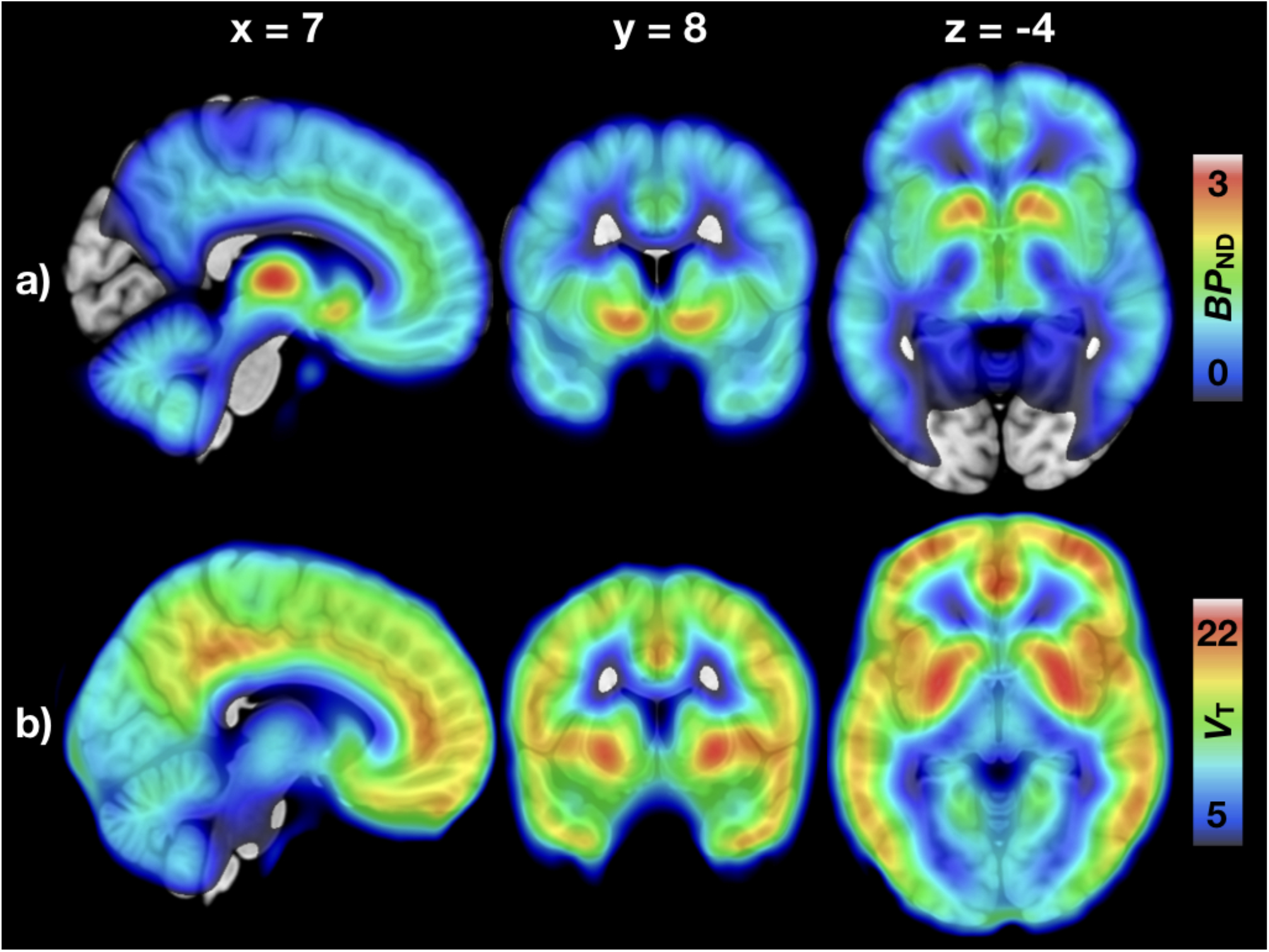
Mean distribution of central µ-opioid and CB_1_-receptors. a) Mean binding potential (*BP*_ND_) of the 92 subjects (70 males and 22 females) studied with [^11^C]carfentanil. b) Mean volume of distribution (*V*_T_) of the 35 males studied with [^18^F]FMPEP-*d*_*2*_.

### 3.1. Regional analysis of neuroreceptor availability and eating behavior

Higher External eating score was associated with lower [^11^C]carfentanil *BP*_ND_ in all *a priori* ROIs (**Figure 2**). In the [^11^C]carfentanil models with other DEBQ subscales and Total DEBQ, the 80% confidence intervals overlapped with zero. For [^18^F]FMPEP-*d*_*2*_, higher Total DEBQ score was associated with lower *V*_T_ all examined ROIs (**Figure 2**). The association directions between *V*_T_ and all DEBQ subscales were negative, but the 95% confidence intervals overlapped with zero. Visualization of the regional relationships between DEBQ scores and neuroreceptor availability in representative ROIs is presented in **Figure 3**.

**Figure 2.**
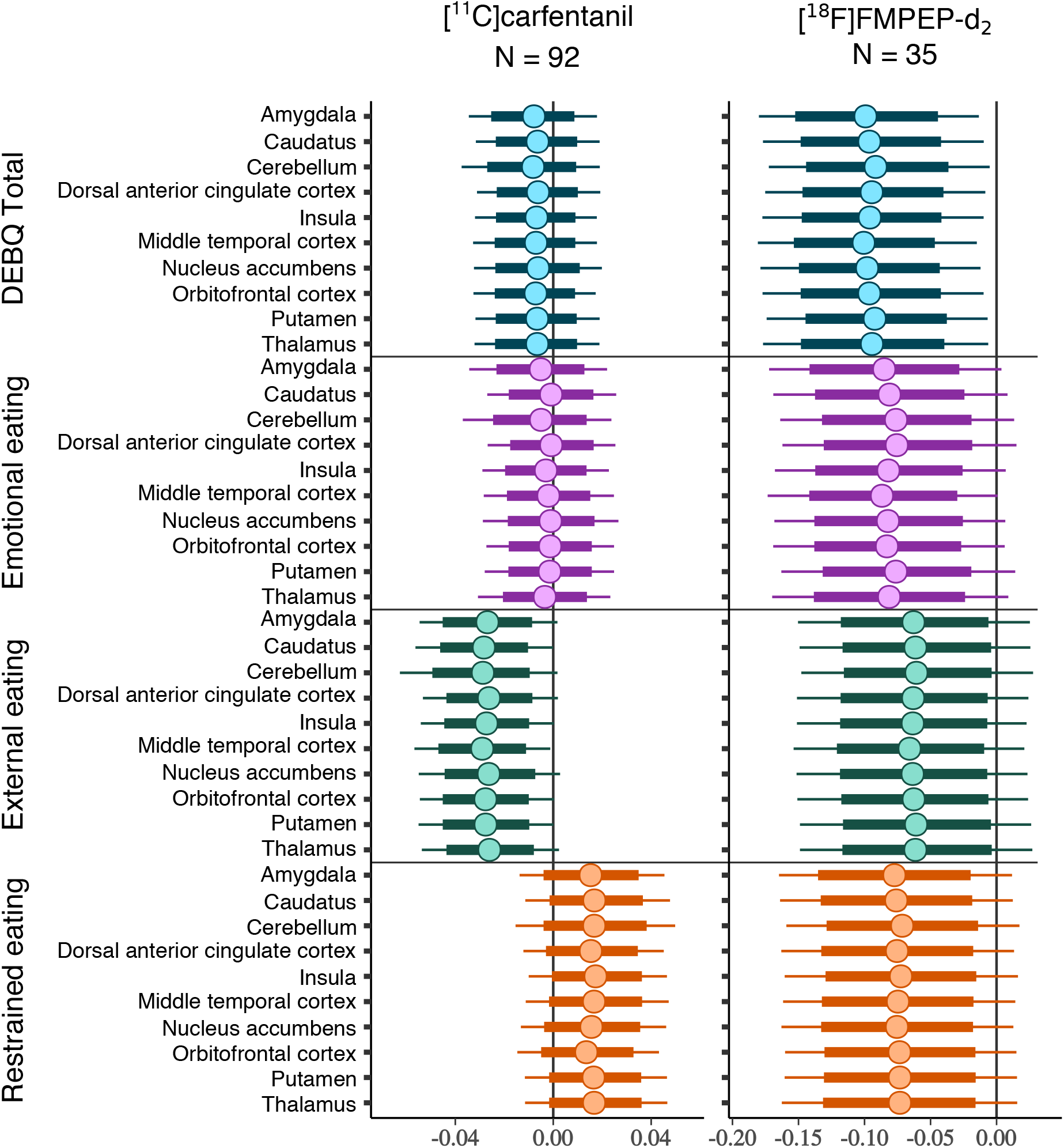
Regional associations of Total DEBQ and subscale scores with µ-opioid and CB_1_-receptor availabilities. The figure shows posterior distributions of the regression coefficients for Total DEBQ and subscale scores on log-transformed [^11^C]carfentanil binding potential (*BP*_ND_) and [^18^F]FMPEP-*d*_*2*_ volume of distribution (*V*_T_) in 10 regions of interest. Age (and PET scanner for [^11^C]carfentanil data) are controlled as covariates. The colored circles represent posterior means, the thick horizontal bars 80% posterior intervals, and the thin horizontal bars 95% posterior intervals.

**Figure 3.**
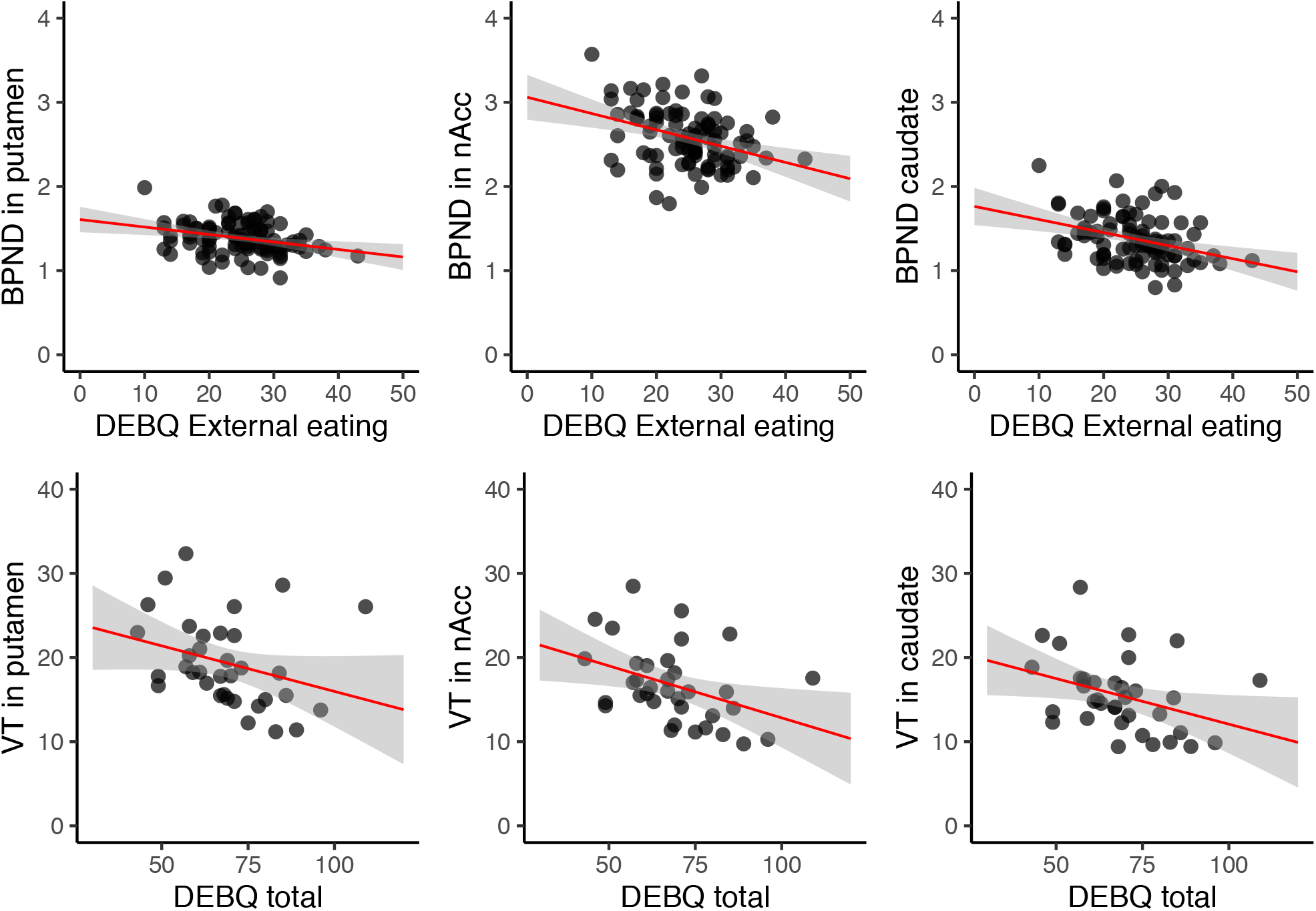
Visualization of regional associations in three representative regions of interest. Upper row: Scatterplots show the associations of External eating score and [^11^C]carfentanil binding potential (*BP*_ND_) in putamen, nucleus accumbens (nAcc) and caudate (92 subjects, LS-regression line with 95% CI). Lower row: Scatterplots show the associations of Total DEBQ score and [^18^F]FMPEP-*d*_*2*_ volume of distribution (*V*_T_) in putamen, nucleus accumbens (nAcc) and caudate (35 subjects, LS-regression line with 95% CI).

### 3.2. Full volume analysis of central receptor availability and eating behavior

For both tracers, full volume results were consistent with the ROI models.

#### 3.2.1. µ-opioid receptor availability and DEBQ

Higher External eating score was associated with lower [^11^C]carfentanil *BP*_ND_ in multiple brain areas (**Figure 4**). Strongest cerebral associations were found in the right frontotemporal cortex and insula (peak voxel coordinates in **Supplementary Table 3**). Associations with Total DEBQ or other subscale scores were not statistically significant. Results were similar in the male subsample (n = 70, **Supplementary Figure 2**) and also when additionally controlling for sex, smoking and BMI, but significant only with more lenient threshold (cluster forming p < 0.05, FWE corrected). In the female subsample, there were no significant associations, likely due to limited statistical power.

**Figure 4.**
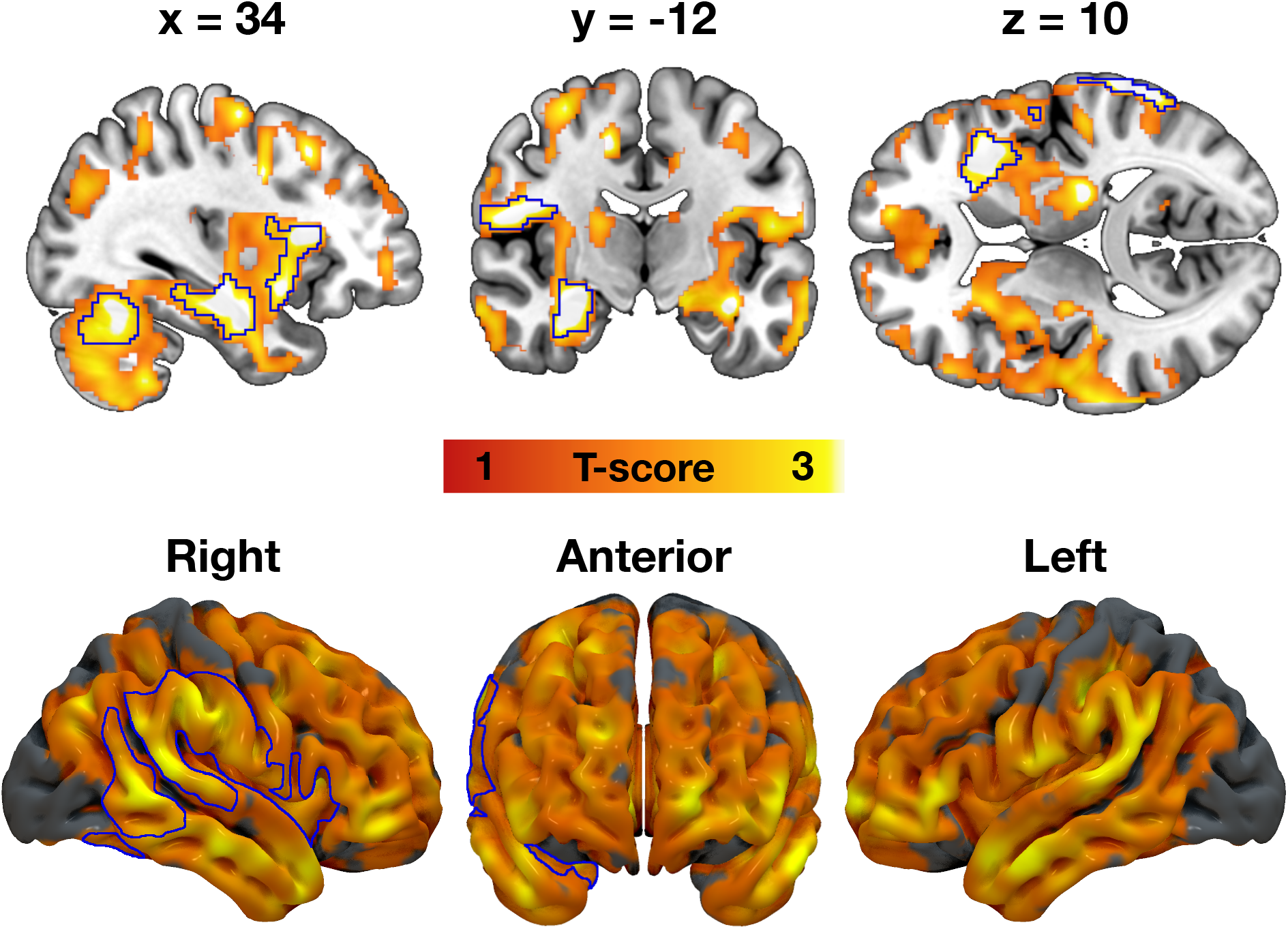
Association between External eating and decreased µ-opioid receptor availability. The blue outline marks brain regions where lower [^11^C]carfentanil binding potential (*BP*_ND_) associated with higher External eating score, age and PET scanner as nuisance covariates, cluster forming threshold p < 0.01, FWE corrected. In the red–yellow T-score scale shown are also additional bilateral associations significant with more lenient cluster-defining threshold (p < 0.05, FWE corrected) for visualization purposes.

#### 3.2.2. CB_1_-receptor availability and DEBQ

Higher Total DEBQ score was associated with lower [^18^F]FMPEP-*d*_*2*_ *V*_T_ bilaterally in multiple brain regions (**Figure 5**). Most prominent associations were found in parahippocampus, frontal striatum, insula, anterior cingulate and frontotemporal cortices (peak voxel coordinates in **Supplementary Table 3**). Full-volume associations with distinct DEBQ subscales and *V*_T_ were not statistically significant. Results were essentially the same when controlling for BMI.

**Figure 5.**
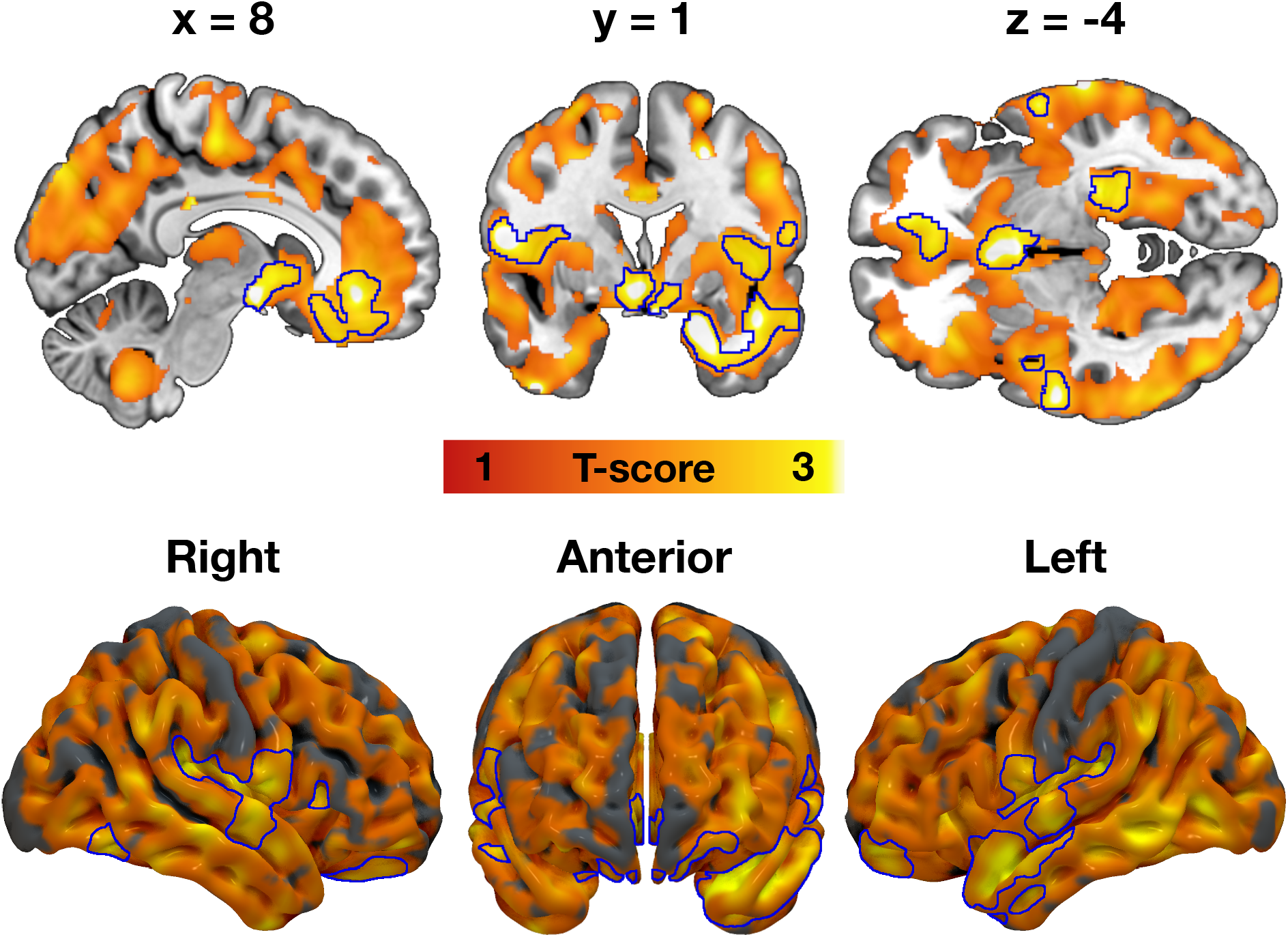
Total Dutch Eating Behavior Questionnaire (DEBQ) score associated with decreased CB_1_-receptor availability. The blue outline marks brain regions where lower [^18^F]FMPEP-*d*_*2*_ volume of distribution (*V*_T_) associated with higher Total DEBQ score, age as a nuisance covariate, cluster forming threshold p < 0.01, FWE corrected. In the red–yellow T-score scale shown are also additional associations significant with more lenient cluster-defining threshold (p < 0.05, FWE corrected) for visualization purposes.

## 4. Discussion

Our main finding was that higher DEBQ scores were associated with lower central availability of µ-opioid and CB_1_-receptors in healthy, non-obese humans. MOR and CB_1_R systems however showed distinct patterns of associations with specific dimensions of self-reported eating: While CB_1_Rs were associated in general negatively with different DEBQ subscale scores (and most saliently with the Total DEBQ score), MORs were specifically and negatively associated with externally driven eating only. Our results support the view that variation in endogenous opioid and endocannabinoid systems explain interindividual variation in feeding, with distinct effects on eating behavior measured with DEBQ.

### 4.1. Central µ-opioid receptors and external eating behavior

External eating – the tendency to feed when encountering palatable food cues such as advertisements – was associated with lowered MOR availability in insula, cortico-limbic regions and striatum, which are major brain areas processing environmental food cues and mediating reward (Berthoud et al., 2017). A bulk of studies have shown that these regions are activated by mere perception of food cues or anticipation of feeding (Nummenmaa et al., 2012; Stice et al., 2008a; Stice et al., 2008b), and our recent work shows that lowered MOR availability is associated with amplified hemodynamic responses to food images in the same regions (Nummenmaa et al., 2018). Higher score on external eating is associated with increased food craving (Burton et al., 2007) and cue-induced palatable food intake (Anschutz et al., 2009; Van Strien et al., 2012a), and may also contribute to short-term weight gain (Van Strien et al., 2012b). Altogether these results suggest that central MOR system has an important role in modulating particularly this kind of impulsive feeding that may lead to overweight.

Previous PET studies have established that feeding triggers endogenous opioid release in humans (Burghardt et al., 2015; Tuulari et al., 2017). Binge eating disorder (BED) is accompanied with downregulated central MORs and high External and Emotional eating scores (Joutsa et al., 2018). Morbid obesity is also associated with lowered central MOR availability (Burghardt et al., 2015; Karlsson et al., 2015), possibly reflecting receptor downregulation due to repeated overstimulation following feeding. In minipigs, already 12 days of high sucrose intake and following central endogenous neurotransmitter release downregulates MORs in cingulate and prefrontal cortices, nucleus accumbens and elsewhere in striatum (Winterdahl et al., 2019). The present findings extend the role of MORs in obesity and eating disorders to different feeding patterns in healthy subjects.

Healthy humans vary considerably in central MOR availability (Kantonen et al., 2020), and it is also possible that lowered MOR availability constitutes a genetically determined (Weerts et al., 2013) risk factor for externally driven eating behavior. In healthy humans, trait impulsivity is associated with central MOR availability (Love et al., 2009). Increased cue-reactivity is prevalent feature of behavioral addictions (Antons et al., 2020), and patients with BED and pathological gambling have reduced availability of central MORs as measured with *in vivo* PET (Majuri et al., 2017). It is thus possible that subjects with lower MOR availability are susceptible for increased external eating in modern environment where they are consistently bombarded with feeding cues in advertisements and food shelves in supermarkets (Berthoud, 2012). However, the present data are purely cross-sectional and longitudinal human studies are needed to further disentangle the causes and the effects between the decrease of MORs in relation to external eating.

### 4.2. Central CB_1_-receptors and eating behavior

Higher Total DEBQ score associated with lower availability of central CB_1_Rs, and ROI-level modeling suggested a consistent negative association with all DEBQ subscales. Compared with the [^11^C]carfentanil model, wider posterior distributions reflect the uncertainty arising from smaller [^18^F]FMPEP-*d*_*2*_ sample size. Brain’s endocannabinoid system is a major homeostatic signaling system, with CB_1_Rs abundant in all known food intake regulating central regions (Bermudez-Silva et al., 2010). In previous brain PET studies, lowered CB_1_R availability has been associated with increased BMI (Hirvonen et al., 2012), while globally upregulated CB_1_Rs have been found in anorexia nervosa (Gérard et al., 2011). These opposite phenotypes on body adiposity spectrum could potentially result from corresponding alterations from CB_1_R-regulated food intake behaviors. Indeed, stimulation of CB_1_Rs by pharmacological agonists or endocannabinoids promotes food intake and amplifies the rewarding properties of feeding (Richard et al., 2009). In contrast, antagonism of the CB_1_Rs by rimonabant (withdrawn anti-obesity drug, Acomplia) effectively reduces food intake and increases energy expenditure, but in many patients also induces psychiatric symptoms (e.g. depressive mood, suicidality, anxiety) (Bermudez-Silva et al., 2010). Accordingly, the endocannabinoid system function has been connected to several other essential behavioral processes in addition to feeding (e.g. stress-coping, emotion regulation, pain perception) (Di Marzo, 2008; Lutz et al., 2015). Being this diverse and complex regulatory system, it may not be possible to pinpoint single distinct aspect of food intake behavior mediated by CB_1_Rs. Rather, our results add support to central CB_1_Rs role in regulation of multiple eating behavior traits, with implications on both homeostatic and hedonic feeding (Quarta et al., 2011).

### 4.3. Limitations and methodological considerations

The [^11^C]carfentanil data were pooled from three PET scanners, which may produce minor variance in outcome measures (Nummenmaa et al., 2020). However, this was accounted for in the analyses by adding the PET scanner as a nuisance covariate to all full-volume and regional analyses. The sample studied with [^11^C]carfentanil consisted predominantly of males, and our statistical power was compromised for detecting potential sex differences. Also all subjects of the [^18^F]FMPEP-*d*_*2*_ subsample were males, and thus conclusions regarding CB_1_Rs might not be generalizable to females. Eating behavior was assessed by self-reports, rather than by direct observations. DEBQ has however been found to successfully identify clinically relevant eating styles (Baños et al., 2014; Wardle, 1987). Our study had a cross-sectional design, and although we found evidence of eating behavior’s association with MOR and CB_1_R systems, whether these receptor systems’ alterations directly promote future weight gain is to be examined in longitudinal studies. Finally, in a single PET scan it is not possible to determine the exact proportions for causal factors to the altered receptor availability, which could potentially be affected by changes in receptor density, affinity or endogenous ligand binding (Henriksen and Willoch, 2008).

## 5. Conclusions

Low cerebral MOR availability is associated with increased externally triggered eating behavior. Modern obesogenic environment may promote food consumption via engaging the opioidergic link of the reward circuit whose tone is linked with cue-reactive eating. Central CB_1_Rs are in turn associated with multiple eating behavioral traits measured with DEBQ, consistent with endocannabinoid system’s role as a major homeostatic regulatory system at the intersection of feeding, emotion and behavior.

## Funding, disclosure and acknowledgements

Declarations of interest: none. This work was supported by Academy of Finland grants (#332225, #304385, and #294897 to LN) and Sigrid Juselius Foundation. The work was also funded by Centre of Excellence of Cardiovascular and Metabolic Diseases supported by Academy of Finland (PN). We are grateful to Emil Aaltonen Foundation, Finnish Cultural Foundation (Southwest Finland Fund), and Jenny and Antti Wihuri Foundation for personal grants to TaK. We thank Turunmaa Duodecim Society, Turku University Hospital Foundation for Education and Research, and Jalmari and Rauha Ahokas Foundation for personal grants to LP. We are grateful to Academy of Finland (grant #256836) and Finnish Alcohol Research Foundation for personal grants to VK. Earlier version of the manuscript has been posted on preprint server bioRxiv (https://www.biorxiv.org/content/10.1101/2020.12.17.423284v1).

## Data and code availability statement

The code for PET data processing pipeline (Magia) is available at https://github.com/tkkarjal/magia.

## Author contributions

TaK: Corresponding and first author, study design, study coordination, data acquisition, data modeling, statistical analysis, interpretation of the results, tables and figures, main writer of the manuscript. ToK: Study design, data modeling, statistical analysis, interpretation of the results, figures, writing of the manuscript. LP: Data acquisition, interpretation of the results, writing of the manuscript. JI: Data acquisition, interpretation of the results, writing of the manuscript. KK: Interpretation of the results, writing of the manuscript. VK: Interpretation of the results, writing of the manuscript. JH: Data modeling, interpretation of the results, writing of the manuscript. PN: Study design, study coordination, interpretation of the results, writing of the manuscript. LN: Study design, study coordination, statistical analysis, interpretation of the results, figures, writing of the manuscript, supervision of the study.

## Supplementary Material

Supplementary Material accompanies this paper.

## References

Anschutz, D.J., Van Strien, T., Van De Ven, M.O.M., Engels, R.C.M.E., 2009. Eating styles and energy intake in young women. Appetite 53, 119–122.

Antons, S., Brand, M., Potenza, M.N., 2020. Neurobiology of cue-reactivity, craving, and inhibitory control in non-substance addictive behaviors. Journal of the Neurological Sciences 415, 116952.

Baños, R.M., Cebolla, A., Moragrega, I., Van Strien, T., Fernández-Aranda, F., Agüera, Z., de la Torre, R., Casanueva, F.F., Fernández-Real, J.M., Fernández-García, J.C., Frühbeck, G., Gómez-Ambrosi, J., Jiménez-Murcia, S., Rodríguez, R., Tinahones, F.J., Botella, C., 2014. Relationship between eating styles and temperament in an Anorexia Nervosa, Healthy Control, and Morbid Obesity female sample. Appetite 76, 76–83.

Barrada, J.R., Van Strien, T., Cebolla, A., 2016. Internal structure and measurement invariance of the Dutch eating behavior questionnaire (DEBQ) in a (nearly) representative Dutch community sample. European Eating Disorders Review 24, 503–509.

Bencherif, B., Guarda, A.S., Colantuoni, C., Ravert, H.T., Dannals, R.F., Frost, J.J., 2005. Regional μ-opioid receptor binding in insular cortex is decreased in bulimia nervosa and correlates inversely with fasting behavior. Journal of Nuclear Medicine 46, 1349–1351.

Bermudez-Silva, F.J., Cardinal, P., Cota, D., 2012. The role of the endocannabinoid system in the neuroendocrine regulation of energy balance. Journal of psychopharmacology 26, 114–124.

Bermudez-Silva, F.J., Viveros, M.P., McPartland, J.M., Rodriguez de Fonseca, F., 2010. The endocannabinoid system, eating behavior and energy homeostasis: The end or a new beginning? Pharmacology Biochemistry and Behavior 95, 375–382.

Berthoud, H.-R., 2012. The neurobiology of food intake in an obesogenic environment. The Proceedings of the Nutrition Society 71, 478–487.

Berthoud, H.-R., Münzberg, H., Morrison, C.D., 2017. Blaming the brain for obesity: integration of hedonic and homeostatic mechanisms. Gastroenterology 152, 1728–1738.

Burghardt, P.R., Rothberg, A.E., Dykhuis, K.E., Burant, C.F., Zubieta, J.-K., 2015. Endogenous Opioid Mechanisms Are Implicated in Obesity and Weight Loss in Humans. The Journal of Clinical Endocrinology & Metabolism 100, 3193–3201.

Burton, P., Smit, H.J., Lightowler, H.J., 2007. The influence of restrained and external eating patterns on overeating. Appetite 49, 191–197.

Castro, D.C., Berridge, K.C., 2017. Opioid and orexin hedonic hotspots in rat orbitofrontal cortex and insula. Proceedings of the National Academy of Sciences 114, E9125–E9134.

Cebolla, A., Barrada, J.R., van Strien, T., Oliver, E., Baños, R., 2014. Validation of the Dutch Eating Behavior Questionnaire (DEBQ) in a sample of Spanish women. Appetite 73, 58–64.

Cota, D., Tschöp, M.H., Horvath, T.L., Levine, A.S., 2006. Cannabinoids, opioids and eating behavior: the molecular face of hedonism? Brain research reviews 51, 85–107.

Di Marzo, V., 2008. Targeting the endocannabinoid system: to enhance or reduce? Nature Reviews Drug Discovery 7, 438–455.

Di Marzo, V., Després, J.-P., 2009. CB1 antagonists for obesity—what lessons have we learned from rimonabant? Nature Reviews Endocrinology 5, 633–638.

French, S.A., Epstein, L.H., Jeffery, R.W., Blundell, J.E., Wardle, J., 2012. Eating behavior dimensions. Associations with energy intake and body weight. A review. Appetite 59, 541–549.

Frost, J.J., Douglass, K.H., Mayberg, H.S., Dannals, R.F., Links, J.M., Wilson, A.A., Ravert, H.T., Crozier, W.C., Wagner, H.N., 1989. Multicompartmental Analysis of [11C]-Carfentanil Binding to Opiate Receptors in Humans Measured by Positron Emission Tomography. Journal of Cerebral Blood Flow & Metabolism 9, 398–409.

Gelman, A., Hill, J., Yajima, M., 2012. Why we (usually) don’t have to worry about multiple comparisons. Journal of Research on Educational Effectiveness 5, 189–211.

Gérard, N., Pieters, G., Goffin, K., Bormans, G., Van Laere, K., 2011. Brain Type 1 Cannabinoid Receptor Availability in Patients with Anorexia and Bulimia Nervosa. Biological Psychiatry 70, 777–784.

Gosnell, B.A., Levine, A.S., 2009. Reward systems and food intake: role of opioids. Int J Obes (Lond) 33 Suppl 2, S54–58.

Guyenet, S.J., Schwartz, M.W., 2012. Regulation of Food Intake, Energy Balance, and Body Fat Mass: Implications for the Pathogenesis and Treatment of Obesity. The Journal of Clinical Endocrinology & Metabolism 97, 745–755.

Henriksen, G., Willoch, F., 2008. Imaging of opioid receptors in the central nervous system. Brain 131, 1171–1196.

Hill, J.O., Peters, J.C., 1998. Environmental contributions to the obesity epidemic. Science 280, 1371–1374.

Hirvonen, J., Goodwin, R.S., Li, C.T., Terry, G.E., Zoghbi, S.S., Morse, C., Pike, V.W., Volkow, N.D., Huestis, M.A., Innis, R.B., 2012. Reversible and regionally selective downregulation of brain cannabinoid CB1 receptors in chronic daily cannabis smokers. Mol Psychiatry 17, 642–649.

Innis, R.B., Cunningham, V.J., Delforge, J., Fujita, M., Gjedde, A., Gunn, R.N., Holden, J., Houle, S., Huang, S.C., Ichise, M., Iida, H., Ito, H., Kimura, Y., Koeppe, R.A., Knudsen, G.M., Knuuti, J., Lammertsma, A.A., Laruelle, M., Logan, J., Maguire, R.P., Mintun, M.A., Morris, E.D., Parsey, R., Price, J.C., Slifstein, M., Sossi, V., Suhara, T., Votaw, J.R., Wong, D.F., Carson, R.E., 2007. Consensus nomenclature for in vivo imaging of reversibly binding radioligands. J Cereb Blood Flow Metab 27, 1533–1539.

Joutsa, J., Karlsson, H.K., Majuri, J., Nuutila, P., Helin, S., Kaasinen, V., Nummenmaa, L., 2018. Binge eating disorder and morbid obesity are associated with lowered mu-opioid receptor availability in the brain. Psychiatry Research: Neuroimaging 276, 41–45.

Kantonen, T., Karjalainen, T., Isojärvi, J., Nuutila, P., Tuisku, J., Rinne, J., Hietala, J., Kaasinen, V., Kalliokoski, K., Scheinin, H., Hirvonen, J., Vehtari, A., Nummenmaa, L., 2020. Interindividual variability and lateralization of μ-opioid receptors in the human brain. Neuroimage 217, 116922.

Karjalainen, T., Tuisku, J., Santavirta, S., Kantonen, T., Bucci, M., Tuominen, L., Hirvonen, J., Hietala, J., Rinne, J.O., Nummenmaa, L., 2020. Magia: Robust Automated Image Processing and Kinetic Modeling Toolbox for PET Neuroinformatics. Frontiers in Neuroinformatics 14.

Karlsson, H.K., Tuominen, L., Tuulari, J.J., Hirvonen, J., Parkkola, R., Helin, S., Salminen, P., Nuutila, P., Nummenmaa, L., 2015. Obesity is associated with decreased mu-opioid but unaltered dopamine D2 receptor availability in the brain. J Neurosci 35, 3959–3965.

Lahesmaa, M., Eriksson, O., Gnad, T., Oikonen, V., Bucci, M., Hirvonen, J., Koskensalo, K., Teuho, J., Niemi, T., Taittonen, M., 2018. Cannabinoid type 1 receptors are upregulated during acute activation of brown adipose tissue. Diabetes 67, 1226–1236.

Larson, N., Story, M., 2009. A review of environmental influences on food choices. Annals of Behavioral Medicine 38, 56–73.

Locke, A.E., Kahali, B., Berndt, S.I., Justice, A.E., Pers, T.H., Day, F.R., Powell, C., Vedantam, S., Buchkovich, M.L., Yang, J., Croteau-Chonka, D.C., Esko, T., Fall, T., Ferreira, T., Gustafsson, S., Kutalik, Z., Luan, J.a., Mägi, R., Randall, J.C., Winkler, T.W., Wood, A.R., Workalemahu, T., Faul, J.D., Smith, J.A., Zhao, J.H., Zhao, W., Chen, J., Fehrmann, R., Hedman, Å.K., Karjalainen, J., Schmidt, E.M., Absher, D., Amin, N., Anderson, D., Beekman, M., Bolton, J.L., Bragg-Gresham, J.L., Buyske, S., Demirkan, A., Deng, G., Ehret, G.B., Feenstra, B., Feitosa, M.F., Fischer, K., Goel, A., Gong, J., Jackson, A.U., Kanoni, S., Kleber, M.E., Kristiansson, K., Lim, U., Lotay, V., Mangino, M., Leach, I.M., Medina-Gomez, C., Medland, S.E., Nalls, M.A., Palmer, C.D., Pasko, D., Pechlivanis, S., Peters, M.J., Prokopenko, I., Shungin, D., Stančáková, A., Strawbridge, R.J., Sung, Y.J., Tanaka, T., Teumer, A., Trompet, S., van der Laan, S.W., van Setten, J., Van Vliet-Ostaptchouk, J.V., Wang, Z., Yengo, L., Zhang, W., Isaacs, A., Albrecht, E., Ärnlöv, J., Arscott, G.M., Attwood, A.P., Bandinelli, S., Barrett, A., Bas, I.N., Bellis, C., Bennett, A.J., Berne, C., Blagieva, R., Blüher, M., Böhringer, S., Bonnycastle, L.L., Böttcher, Y., Boyd, H.A., Bruinenberg, M., Caspersen, I.H., Chen, Y.-D.I., Clarke, R., Daw, E.W., de Craen, A.J.M., Delgado, G., Dimitriou, M., Doney, A.S.F., Eklund, N., Estrada, K., Eury, E., Folkersen, L., Fraser, R.M., Garcia, M.E., Geller, F., Giedraitis, V., Gigante, B., Go, A.S., Golay, A., Goodall, A.H., Gordon, S.D., Gorski, M., Grabe, H.-J., Grallert, H., Grammer, T.B., Gräßler, J., Grönberg, H., Groves, C.J., Gusto, G., Haessler, J., Hall, P., Haller, T., Hallmans, G., Hartman, C.A., Hassinen, M., Hayward, C., Heard-Costa, N.L., Helmer, Q., Hengstenberg, C., Holmen, O., Hottenga, J.-J., James, A.L., Jeff, J.M., Johansson, Å., Jolley, J., Juliusdottir, T., Kinnunen, L., Koenig, W., Koskenvuo, M., Kratzer, W., Laitinen, J., Lamina, C., Leander, K., Lee, N.R., Lichtner, P., Lind, L., Lindström, J., Lo, K.S., Lobbens, S., Lorbeer, R., Lu, Y., Mach, F., Magnusson, P.K.E., Mahajan, A., McArdle, W.L., McLachlan, S., Menni, C., Merger, S., Mihailov, E., Milani, L., Moayyeri, A., Monda, K.L., Morken, M.A., Mulas, A., Müller, G., Müller-Nurasyid, M., Musk, A.W., Nagaraja, R., Nöthen, M.M., Nolte, I.M., Pilz, S., Rayner, N.W., Renstrom, F., Rettig, R., Ried, J.S., Ripke, S., Robertson, N.R., Rose, L.M., Sanna, S., Scharnagl, H., Scholtens, S., Schumacher, F.R., Scott, W.R., Seufferlein, T., Shi, J., Smith, A.V., Smolonska, J., Stanton, A.V., Steinthorsdottir, V., Stirrups, K., Stringham, H.M., Sundström, J., Swertz, M.A., Swift, A.J., Syvänen, A.-C., Tan, S.-T., Tayo, B.O., Thorand, B., Thorleifsson, G., Tyrer, J.P., Uh, H.-W., Vandenput, L., Verhulst, F.C., Vermeulen, S.H., Verweij, N., Vonk, J.M., Waite, L.L., Warren, H.R., Waterworth, D., Weedon, M.N., Wilkens, L.R., Willenborg, C., Wilsgaard, T., Wojczynski, M.K., Wong, A., Wright, A.F., Zhang, Q., LifeLines Cohort, S., Brennan, E.P., Choi, M., Dastani, Z., Drong, A.W., Eriksson, P., Franco-Cereceda, A., Gådin, J.R., Gharavi, A.G., Goddard, M.E., Handsaker, R.E., Huang, J., Karpe, F., Kathiresan, S., Keildson, S., Kiryluk, K., Kubo, M., Lee, J.-Y., Liang, L., Lifton, R.P., Ma, B., McCarroll, S.A., McKnight, A.J., Min, J.L., Moffatt, M.F., Montgomery, G.W., Murabito, J.M., Nicholson, G., Nyholt, D.R., Okada, Y., Perry, J.R.B., Dorajoo, R., Reinmaa, E., Salem, R.M., Sandholm, N., Scott, R.A., Stolk, L., Takahashi, A., Tanaka, T., van ‘t Hooft, F.M., Vinkhuyzen, A.A.E., Westra, H.-J., Zheng, W., Zondervan, K.T., Consortium, A.D., Group, A.-B.W., Consortium, C.A.D., Consortium, C.K., Glgc, Icbp, Investigators, M., Mu, T.C., Consortium, M.I., Consortium, P., ReproGen, C., Consortium, G., International Endogene, C., Heath, A.C., Arveiler, D., Bakker, S.J.L., Beilby, J., Bergman, R.N., Blangero, J., Bovet, P., Campbell, H., Caulfield, M.J., Cesana, G., Chakravarti, A., Chasman, D.I., Chines, P.S., Collins, F.S., Crawford, D.C., Cupples, L.A., Cusi, D., Danesh, J., de Faire, U., den Ruijter, H.M., Dominiczak, A.F., Erbel, R., Erdmann, J., Eriksson, J.G., Farrall, M., Felix, S.B., Ferrannini, E., Ferrières, J., Ford, I., Forouhi, N.G., Forrester, T., Franco, O.H., Gansevoort, R.T., Gejman, P.V., Gieger, C., Gottesman, O., Gudnason, V., Gyllensten, U., Hall, A.S., Harris, T.B., Hattersley, A.T., Hicks, A.A., Hindorff, L.A., Hingorani, A.D., Hofman, A., Homuth, G., Hovingh, G.K., Humphries, S.E., Hunt, S.C., Hyppönen, E., Illig, T., Jacobs, K.B., Jarvelin, M.-R., Jöckel, K.-H., Johansen, B., Jousilahti, P., Jukema, J.W., Jula, A.M., Kaprio, J., Kastelein, J.J.P., Keinanen-Kiukaanniemi, S.M., Kiemeney, L.A., Knekt, P., Kooner, J.S., Kooperberg, C., Kovacs, P., Kraja, A.T., Kumari, M., Kuusisto, J., Lakka, T.A., Langenberg, C., Marchand, L.L., Lehtimäki, T., Lyssenko, V., Männistö, S., Marette, A., Matise, T.C., McKenzie, C.A., McKnight, B., Moll, F.L., Morris, A.D., Morris, A.P., Murray, J.C., Nelis, M., Ohlsson, C., Oldehinkel, A.J., Ong, K.K., Madden, P.A.F., Pasterkamp, G., Peden, J.F., Peters, A., Postma, D.S., Pramstaller, P.P., Price, J.F., Qi, L., Raitakari, O.T., Rankinen, T., Rao, D.C., Rice, T.K., Ridker, P.M., Rioux, J.D., Ritchie, M.D., Rudan, I., Salomaa, V., Samani, N.J., Saramies, J., Sarzynski, M.A., Schunkert, H., Schwarz, P.E.H., Sever, P., Shuldiner, A.R., Sinisalo, J., Stolk, R.P., Strauch, K., Tönjes, A., Trégouët, D.-A., Tremblay, A., Tremoli, E., Virtamo, J., Vohl, M.-C., Völker, U., Waeber, G., Willemsen, G., Witteman, J.C., Zillikens, M.C., Adair, L.S., Amouyel, P., Asselbergs, F.W., Assimes, T.L., Bochud, M., Boehm, B.O., Boerwinkle, E., Bornstein, S.R., Bottinger, E.P., Bouchard, C., Cauchi, S., Chambers, J.C., Chanock, S.J., Cooper, R.S., de Bakker, P.I.W., Dedoussis, G., Ferrucci, L., Franks, P.W., Froguel, P., Groop, L.C., Haiman, C.A., Hamsten, A., Hui, J., Hunter, D.J., Hveem, K., Kaplan, R.C., Kivimaki, M., Kuh, D., Laakso, M., Liu, Y., Martin, N.G., März, W., Melbye, M., Metspalu, A., Moebus, S., Munroe, P.B., Njølstad, I., Oostra, B.A., Palmer, C.N.A., Pedersen, N.L., Perola, M., Pérusse, L., Peters, U., Power, C., Quertermous, T., Rauramaa, R., Rivadeneira, F., Saaristo, T.E., Saleheen, D., Sattar, N., Schadt, E.E., Schlessinger, D., Slagboom, P.E., Snieder, H., Spector, T.D., Thorsteinsdottir, U., Stumvoll, M., Tuomilehto, J., Uitterlinden, A.G., Uusitupa, M., van der Harst, P., Walker, M., Wallaschofski, H., Wareham, N.J., Watkins, H., Weir, D.R., Wichmann, H.E., Wilson, J.F., Zanen, P., Borecki, I.B., Deloukas, P., Fox, C.S., Heid, I.M., O’Connell, J.R., Strachan, D.P., Stefansson, K., van Duijn, C.M., Abecasis, G.R., Franke, L., Frayling, T.M., McCarthy, M.I., Visscher, P.M., Scherag, A., Willer, C.J., Boehnke, M., Mohlke, K.L., Lindgren, C.M., Beckmann, J.S., Barroso, I., North, K.E., Ingelsson, E., Hirschhorn, J.N., Loos, R.J.F., Speliotes, E.K., 2015. Genetic studies of body mass index yield new insights for obesity biology. Nature 518, 197–206.

Logan, J., 2000. Graphical analysis of PET data applied to reversible and irreversible tracers. Nuclear Medicine and Biology 27, 661–670.

Love, T.M., Stohler, C.S., Zubieta, J.-K., 2009. Positron emission tomography measures of endogenous opioid neurotransmission and impulsiveness traits in humans. Arch Gen Psychiatry 66, 1124–1134.

Lutz, B., Marsicano, G., Maldonado, R., Hillard, C.J., 2015. The endocannabinoid system in guarding against fear, anxiety and stress. Nature Reviews Neuroscience 16, 705–718.

Majuri, J., Joutsa, J., Johansson, J., Voon, V., Alakurtti, K., Parkkola, R., Lahti, T., Alho, H., Hirvonen, J., Arponen, E., Forsback, S., Kaasinen, V., 2017. Dopamine and Opioid Neurotransmission in Behavioral Addictions: A Comparative PET Study in Pathological Gambling and Binge Eating. Neuropsychopharmacology 42, 1169–1177.

Malesza, M., Kaczmarek, M.C., 2019. One year reliability of the Dutch eating behavior questionnaire: an extension into clinical population. Journal of Public Health, 1–7.

Mechoulam, R., Parker, L.A., 2013. The endocannabinoid system and the brain. Annu Rev Psychol 64, 21–47.

Mena, J.D., Sadeghian, K., Baldo, B.A., 2011. Induction of hyperphagia and carbohydrate intake by μ-opioid receptor stimulation in circumscribed regions of frontal cortex. Journal of Neuroscience 31, 3249–3260.

Nogueiras, R., Romero-Pico, A., Vazquez, M.J., Novelle, M.G., Lopez, M., Dieguez, C., 2012. The opioid system and food intake: homeostatic and hedonic mechanisms. Obes Facts 5, 196–207.

Nummenmaa, L., Hirvonen, J., Hannukainen, J.C., Immonen, H., Lindroos, M.M., Salminen, P., Nuutila, P., 2012. Dorsal striatum and its limbic connectivity mediate abnormal anticipatory reward processing in obesity. PLoS One 7, e31089.

Nummenmaa, L., Karjalainen, T., Isojärvi, J., Kantonen, T., Tuisku, J., Kaasinen, V., Joutsa, J., Nuutila, P., Kalliokoski, K., Hirvonen, J., Hietala, J., Rinne, J., 2020. Lowered endogenous muopioid receptor availability in subclinical depression and anxiety. Neuropsychopharmacology.

Nummenmaa, L., Saanijoki, T., Tuominen, L., Hirvonen, J., Tuulari, J.J., Nuutila, P., Kalliokoski, K., 2018. μ-opioid receptor system mediates reward processing in humans. Nature Communications 9, 1500.

Quarta, C., Mazza, R., Obici, S., Pasquali, R., Pagotto, U., 2011. Energy balance regulation by endocannabinoids at central and peripheral levels. Trends in Molecular Medicine 17, 518–526.

Richard, D., Guesdon, B., Timofeeva, E., 2009. The brain endocannabinoid system in the regulation of energy balance. Best Practice & Research Clinical Endocrinology & Metabolism 23, 17–32.

Rowland, N.E., Mukherjee, M., Robertson, K., 2001. Effects of the cannabinoid receptor antagonist SR 141716, alone and in combination with dexfenfluramine or naloxone, on food intake in rats. Psychopharmacology 159, 111–116.

Silventoinen, K., Konttinen, H., 2020. Obesity and eating behavior from the perspective of twin and genetic research. Neuroscience & biobehavioral reviews 109, 150–165.

Smith, K.S., Berridge, K.C., 2007. Opioid limbic circuit for reward: interaction between hedonic hotspots of nucleus accumbens and ventral pallidum. Journal of Neuroscience 27, 1594–1605.

Solinas, M., Goldberg, S.R., 2005. Motivational Effects of Cannabinoids and Opioids on Food Reinforcement Depend on Simultaneous Activation of Cannabinoid and Opioid Systems. Neuropsychopharmacology 30, 2035–2045.

Solinas, M., Zangen, A., Thiriet, N., Goldberg, S.R., 2004. Beta-endorphin elevations in the ventral tegmental area regulate the discriminative effects of Delta-9-tetrahydrocannabinol. Eur J Neurosci 19, 3183–3192.

Stice, E., Spoor, S., Bohon, C., Small, D., 2008a. Relation between obesity and blunted striatal response to food is moderated by TaqIA A1 allele. Science 322, 449–452.

Stice, E., Spoor, S., Bohon, C., Veldhuizen, M.G., Small, D.M., 2008b. Relation of reward from food intake and anticipated food intake to obesity: a functional magnetic resonance imaging study. Journal of abnormal psychology 117, 924.

Terry, G.E., Hirvonen, J., Liow, J.-S., Zoghbi, S.S., Gladding, R., Tauscher, J.T., Schaus, J.M., Phebus, L., Felder, C.C., Morse, C.L., 2010. Imaging and quantitation of cannabinoid CB1 receptors in human and monkey brains using 18F-labeled inverse agonist radioligands. Journal of Nuclear Medicine 51, 112–120.

Tuulari, J.J., Tuominen, L., de Boer, F.E., Hirvonen, J., Helin, S., Nuutila, P., Nummenmaa, L., 2017. Feeding Releases Endogenous Opioids in Humans. J Neurosci 37, 8284–8291.

Van Strien, T., Frijters, J.E., Bergers, G.P., Defares, P.B., 1986a. The Dutch Eating Behavior Questionnaire (DEBQ) for assessment of restrained, emotional, and external eating behavior. International journal of eating disorders 5, 295–315.

Van Strien, T., Frijters, J.E., Van Staveren, W.A., Defares, P.B., Deurenberg, P., 1986b. The predictive validity of the Dutch restrained eating scale. International journal of eating disorders 5, 747–755.

Van Strien, T., Herman, C.P., Verheijden, M.W., 2009. Eating style, overeating, and overweight in a representative Dutch sample. Does external eating play a role? Appetite 52, 380–387.

Van Strien, T., Peter Herman, C., Anschutz, D., 2012a. The predictive validity of the DEBQ-external eating scale for eating in response to food commercials while watching television. International journal of eating disorders 45, 257–262.

Van Strien, T., Peter Herman, C., Verheijden, M.W., 2012b. Eating style, overeating and weight gain. A prospective 2-year follow-up study in a representative Dutch sample. Appetite 59, 782–789.

Van Strien, T., Schippers, G.M., Cox, W.M., 1995. On the relationship between emotional and external eating behavior. Addictive Behaviors 20, 585–594.

Vigano, D., Valenti, M., Cascio, M.G., Di Marzo, V., Parolaro, D., Rubino, T., 2004. Changes in endocannabinoid levels in a rat model of behavioural sensitization to morphine. Eur J Neurosci 20, 1849–1857.

Wardle, J., 1987. Eating style: a validation study of the Dutch Eating Behaviour Questionnaire in normal subjects and women with eating disorders. Journal of psychosomatic research 31, 161–169.

Weerts, E.M., McCaul, M.E., Kuwabara, H., Yang, X., Xu, X., Dannals, R.F., Frost, J.J., Wong, D.F., Wand, G.S., 2013. Influence of OPRM1 Asn40Asp variant (A118G) on [11C]carfentanil binding potential: preliminary findings in human subjects. Int J Neuropsychopharmacol 16, 47–53.

Winterdahl, M., Noer, O., Orlowski, D., Schacht, A.C., Jakobsen, S., Alstrup, A.K., Gjedde, A., Landau, A.M., 2019. Sucrose intake lowers μ-opioid and dopamine D2/3 receptor availability in porcine brain. Scientific reports 9, 1–11.

Yeomans, M.R., Gray, R.W., 2002. Opioid peptides and the control of human ingestive behaviour. Neurosci Biobehav Rev 26, 713–728.

Yeomans, M.R., Leitch, M., Mobini, S., 2008. Impulsivity is associated with the disinhibition but not restraint factor from the Three Factor Eating Questionnaire. Appetite 50, 469–476.

Ziauddeen, H., Chamberlain, S.R., Nathan, P.J., Koch, A., Maltby, K., Bush, M., Tao, W.X., Napolitano, A., Skeggs, A.L., Brooke, A.C., Cheke, L., Clayton, N.S., Sadaf Farooqi, I., O’Rahilly, S., Waterworth, D., Song, K., Hosking, L., Richards, D.B., Fletcher, P.C., Bullmore, E.T., 2013. Effects of the mu-opioid receptor antagonist GSK1521498 on hedonic and consummatory eating behaviour: a proof of mechanism study in binge-eating obese subjects. Mol Psychiatry 18, 1287–1293.

Zubieta, J.K., Dannals, R.F., Frost, J.J., 1999. Gender and age influences on human brain mu-opioid receptor binding measured by PET. Am J Psychiatry 156, 842–848.

