## Supplementary Material for "Cerebral μ-opioid and CB_1_-receptor systems have distinct roles in human feeding behavior"

i. PET scanners and smoking status

ii. Descriptive correlations of the sample

iii. Table of the p-values for descriptive correlations

iv. Association between External eating score and μ-opioid receptor availability in subsamples of 70 males and 22 females

v. Peak voxel coordinates for the full volume associations

**PET scanners and smoking status**

**Supplementary Table 1**. The information about PET scanners and smoking status of the sample.

| | Males (n = 70) | Females (n = 22) |
|---|---|---|
| PET scanner (n) | | |
| HRRT (Siemens Medical Solutions) | 8 | 9 |
| Discovery 690 PET/CT (GE Healthcare) | 10 | 13 |
| GE Discovery VCT PET/CT (GE Healthcare) | 52 | 0 |
| Smoking status (n)* | | |
| Smoking | 0 | 7 |
| Nonsmoking | 57 | 15 |
| Unknown | 13 | 0 |

* The 13 individuals whose smoking status was unknown were classified as nonsmokers because in Finland, less than 20 % of the adult population smoked cigarettes in 2000 (http://urn.fi/URN:NBN:fi-fe2018102938947). Results were similar in a parallel analysis where individuals with unknown smoking status were classified as smokers.

**Descriptive correlations of the sample**

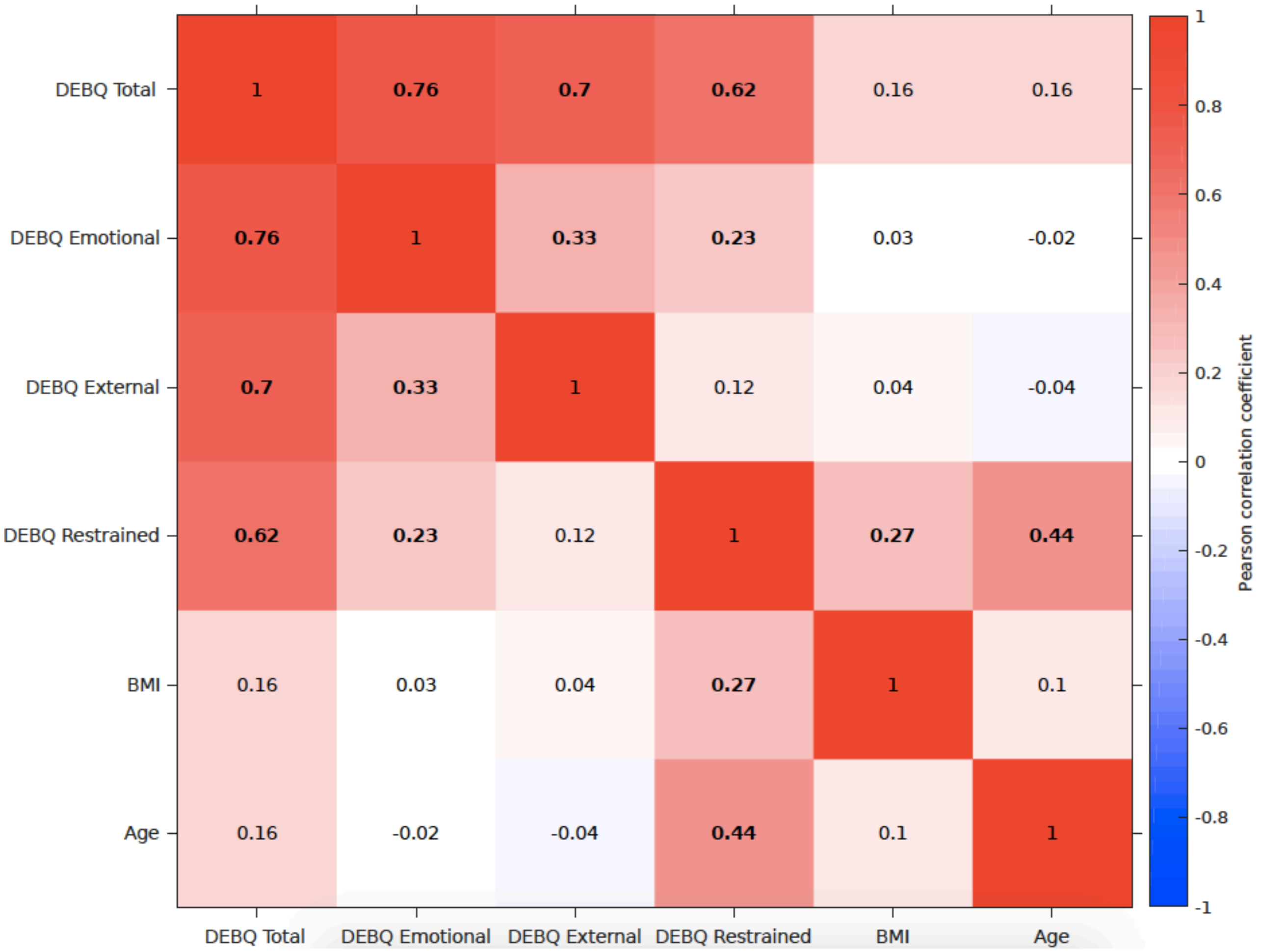


**Supplementary Figure 1**. Descriptive correlations between the Dutch Eating Behaviour Questionnaire (DEBQ) total and subscale scores, age and body mass index (BMI) in the sample of 92 subjects (70 males and 22 females) scanned with [$^{11}$C]carfentanil. The numbers indicate Pearson correlation coefficients. $p < 0.05$ correlations are marked with boldface.

**Table of the p-values for descriptive correlations**

**Supplementary Table 2**. The p-values for corresponding Pearson correlation coefficients in the sample of 92 subjects (70 males and 22 females) scanned with [$^{11}$C]carfentanil.

| | DEBQ Total | DEBQ Emotional | DEBQ External | DEBQ Restrained | BMI | Age |
|---|---|---|---|---|---|---|
| **DEBQ Total** | 1.000 | <0.001 | <0.001 | <0.001 | 0.137 | 0.119 |
| **DEBQ Emotional** | <0.001 | 1.000 | 0.001 | 0.028 | 0.753 | 0.822 |
| **DEBQ External** | <0.001 | 0.001 | 1.000 | 0.267 | 0.728 | 0.676 |
| **DEBQ Restrained** | <0.001 | 0.028 | 0.267 | 1.000 | 0.009 | <0.001 |
| **BMI** | 0.137 | 0.753 | 0.728 | 0.009 | 1.000 | 0.352 |
| **Age** | 0.119 | 0.822 | 0.676 | <0.001 | 0.352 | 1.000 |

**Association between External eating score and μ-opioid receptor availability in subsamples of 70 males and 22 females**

The male subsample

In the subsample of 70 males, the associations between External eating score and μ-opioid receptor availability were mostly similar than with the full sample, associations in caudatus being slightly less prominent.

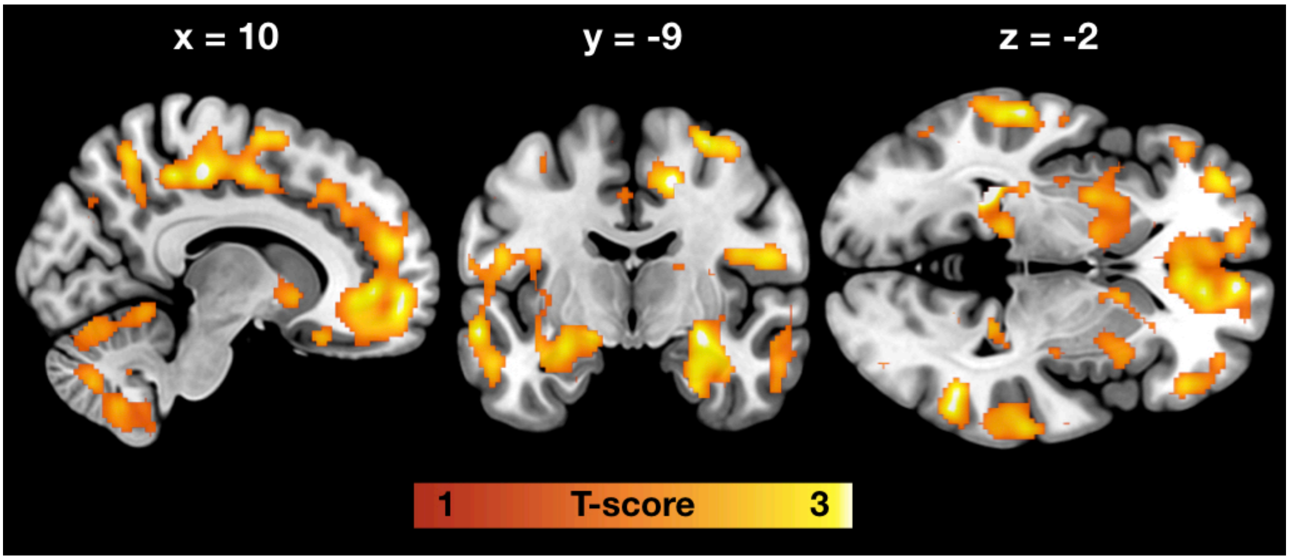


**Supplementary Figure 2**. Brain regions where higher External eating score associated with lower [$^{11}$C]carfentanil binding potential ($BP_{ND}$) in 70 males, age and PET scanner as nuisance covariates. Shown are clusters where $p < 0.05$, FWE corrected at cluster level.

The female subsample

In the subsample of 22 females, there were no statistically significant associations between External eating score and central μ-opioid receptor availability. This is potentially due to the relatively low number of female subjects.

### Peak voxel coordinates for the full volume associations

**Supplementary Table 3**. Peak voxel coordinates for the full volume associations.

**Negative association between [$^{11}$C]carfentanil $BP_{ND}$ and External eating score (n = 92)**

| Brain region | MNI coordinates x | y | z | T-score |
|---|---|---|---|---|
| Right middle temporal gyrus | 66 | -48 | 6 | 4.32 |
| Right postcentral gyrus | 54 | -10 | 18 | 3.85 |
| Right anterior insula | 34 | 20 | 12 | 3.85 |
| Cerebellum | 26 | -66 | -18 | 3.87 |
| | -16 | -60 | -22 | 3.48 |
| | 26 | -52 | -18 | 3.39 |

**Negative association between [$^{18}$F]FMPEP-$d_2$ $V_T$ and Total DEBQ score (n = 35)**

| Brain region | MNI coordinates x | y | z | T-score |
|---|---|---|---|---|
| Left superior temporal gyrus | -33 | 8 | -33 | 4.00 |
| | -48 | 7 | -19 | 3.70 |
| Left parahippocampus | -21 | 1 | -23 | 3.67 |
| Right parahippocampus | 30 | -35 | -17 | 3.63 |
| Right caudate | 20 | -21 | 23 | 3.08 |
| Right cerebellum | 30 | -32 | -30 | 2.96 |
| Left middle frontal gyrus | -23 | 57 | -16 | 3.42 |
| Anterior cingulate cortex | 2 | 2 | -7 | 3.21 |
| Anterior caudate | -1 | 14 | -3 | 3.18 |
| Right insula | 42 | -33 | 20 | 3.42 |
| Right precentral gyrus | 56 | 0 | 13 | 3.39 |
| Right insula | 37 | -12 | 16 | 3.24 |

Cluster forming threshold $p < 0.01$, FWE corrected at cluster level. Age and PET scanner as covariates in the [$^{11}$C]carfentanil model, age as a covariate in the [$^{18}$F]FMPEP-$d_2$ model.
